# PIVOTAL: Prioritizing variants of uncertain significance with spatial genomic patterns in the 3D proteome

**DOI:** 10.1101/2020.06.04.135103

**Authors:** Siqi Liang, Matthew Mort, Peter D. Stenson, David N. Cooper, Haiyuan Yu

## Abstract

Variants of uncertain significance (VUS) have posed an increasingly prominent challenge to clinicians due to their growing numbers and difficulties in making clinical responses to them. Currently there are no existing methods that leverage the spatial relationship of known disease mutations and genomic properties for prioritizing variants of uncertain significance. More importantly, disease genes often associate with multiple clinically distinct diseases, but none of the existing variant prioritization methods provide clues as to the specific type of disease potentially associated with a given variant. We present PIVOTAL, a spatial neighborhood-based method using three-dimensional structural models of proteins, that significantly improves current variant prioritization tools and identifies potential disease etiology of candidate variants on a proteome scale. Using PIVOTAL, we made pathogenicity predictions for over 140,000 VUS and deployed a web application (http://pivotal.yulab.org) that enables users both to explore these data and to perform custom calculations.

---

Fueled by recent advances in next-generation sequencing technologies, the increasing popularity of genetic testing in clinical settings has given rise to an abundance of variants of uncertain clinical significance (VUS). Under the American College of Medical Genetics and Genomics (ACMG) guidelines^1^, these variants have insufficient or conflicting genetic or functional evidence to support their pathogenicity and hence pose serious difficulties both for patients in comprehending their pathological relevance and for clinicians to in deciding upon an appropriate clinical response^2^. As of October 2019, there were more than 140,000 VUS documented in the ClinVar^3^ database (Supplementary Table 1), over 4 times as many as pathogenic and likely pathogenic mutations combined. Our inability to interpret the phenotypic consequences of this prevailing majority of clinically detected variants represents a critical roadblock to personalized medicine.

Missense variants represent a majority of coding mutations in ClinVar (Supplementary. Fig. 1a). Previous studies have explored the distribution patterns of missense disease mutations on protein structures. For example, structural clustering of missense mutations in the three-dimensional space has been observed for both Mendelian diseases^4,5^ and cancer^6^. Furthermore, a majority of disease genes are pleiotropic and often associate with multiple clinically distinct diseases (Supplementary. Fig. 1b-c). For these pleiotropic genes, missense mutations on the same protein-interacting interface tend to cause the same disease^7^. However, these structural clues have been largely overlooked by current approaches for prioritizing VUS, even though as many as ~90% of proteins (Fig. 1a) and ~60% of amino acid residues (Fig. 1b) have been covered by either crystal structures in the Protein Data Bank^8^ or high-quality homology models. Current variant prioritization approaches primarily comprise two classes: functional experimental assays that specifically assess the impact of variants on certain genes, such as *BRCA1*^9^, *LMNA* and *MYBPC3*^10^, and computational tools including PolyPhen-2^11^, SIFT^12^, PROVEAN^13^, CADD^14^, M-CAP^15^, MetaLR^16^, MetaSVM^16^, REVEL^17^ and VEST^18^ that predict the deleteriousness of variants, and can be applied in a genome-wide manner. Although features derived from protein structures have been exploited in mutation prioritization tools such as PolyPhen-2, there is currently no method that uses as evidence the distribution of known disease mutations and genomic measurements from the structural neighborhood of a variant. More importantly, none of these tools provide insight into which specific disease with a prioritized variant might be associated – this is especially important for variants on pleiotropic genes.

**Figure 1.**
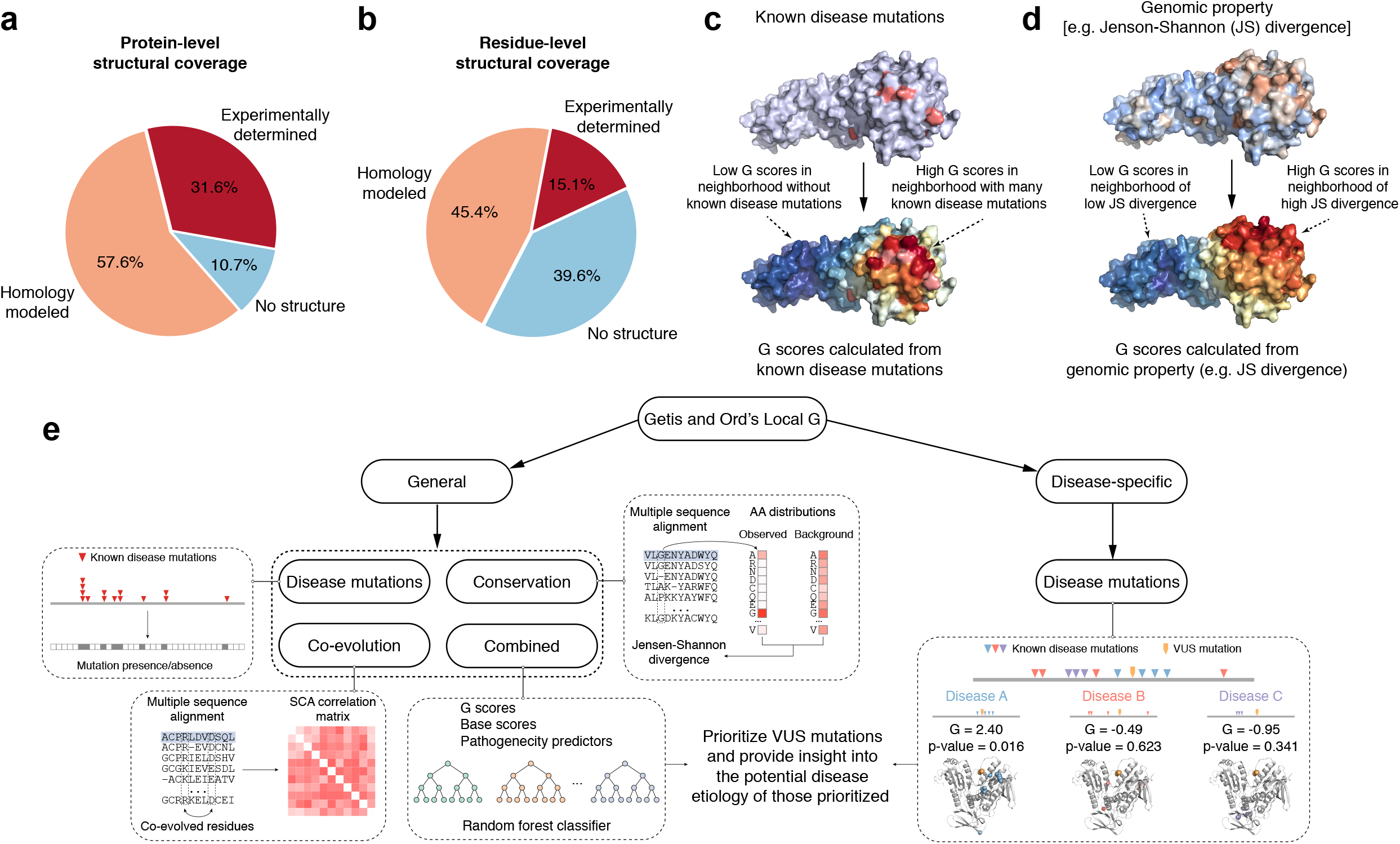
The PIVOTAL framework. (a) Coverage of the human proteome by PDB structures and homology models at the protein level. (b) Coverage of the human proteome by PDB structures and homology models at the amino acid residue level. (c) G score calculated from the presence or abscence of known disease mutations characterizes the extent to which disease mutations cluster in the neighborhood of a given amino acid residue. (d) G score calculated from genomic properties (e.g. Jensen-Shannon divergence) characterizes the extent to which high or low values cluster in the neighborhood of a given amino acid residue. (e) The Getis and Ord’s local G statistic is applied in two major ways: G scores calculated from known disease mutations, evolutionary conservation and co-evolution are combined with existing pathogenicity predictors for VUS prioritization; the disease-specific G score calculated from known pathogenic mutations causing a certain disease provides information about the potential disease association of VUS.

Recently, a number of studies have utilized structural distributional properties for characterizing missense variation. For instance, Gress et al.^19^ and Sivley et al.^20^ have characterized the spatial distribution of different types of missense variant on protein structures, and spatial analysis has been applied to the classification of pathogenic and benign mutations in certain genes^21,22^. In addition, Hicks et al. characterized amino acid residue sites that are intolerant to missense variation by incorporating protein structures and human genetic variation^23^. However, none of these studies have leveraged the abundance of known disease mutations and protein structural information to perform variant prioritization on a proteome scale.

Here, we present PIVOTAL, a framework along with a web server that prioritizes VUS on the entire human structural proteome and provides specific hypotheses of their disease etiology. The distinguishing feature of PIVOTAL is its use of the spatial distribution patterns of known disease mutations and genomic properties on 3D protein structures. This is empowered by an adaptation of Getis and Ord’s local G statistic^24,25^ (see Methods), which characterizes clustering of high or low values in a spatial neighborhood (Fig. 1c-d). Incorporating G statistics calculated from known disease mutations, evolutionary conservation and co-evolution with existing variant deleteriousness predictors enables PIVOTAL to make general pathogenicity predictions for missense mutations. Furthermore, disease-specific G scores calculated from known disease mutations causing the same disease shed light on their potential disease etiology (Fig. 1e).

Using 82,920 known disease mutations in the Human Gene Mutation Database (HGMD)^26^, we calculated G statistics based on the presence or absence of disease mutations on each amino acid residue, and we then evaluated their ability to distinguish 3,706 pathogenic mutations from 6,362 benign ones from ClinVar. To make sure the evaluation is meaningful, we removed all mutations in ClinVar that are on the same amino acid residues as any known disease mutation in HGMD (see Methods). We find that HGMD disease mutation-based G scores are significantly higher for ClinVar pathogenic mutations than for benign mutations (Fig. 2a, P < 10^-20^, Mann-Whitney U test). Furthermore, amino acid residue sites that exhibit significantly high G statistics (see Methods) are enriched for ClinVar pathogenic mutations across different false discovery rates (FDR) (Supplementary Fig. 1d). These results establish disease mutation-derived G scores as an informative feature for pathogenicity prediction. To simulate their use in practice, we compiled five sets of pathogenic and benign mutations across five timestamps benchmarked by ClinVar (Supplementary Table 1). Each time, we used pathogenic mutations in an earlier version and only structural models available at the corresponding timestamp to calculate G statistics; we then evaluated their performance on a later version of pathogenic and benign mutations. We observed significantly higher G scores for pathogenic mutations irrespective of the version combination (Fig. 2b, P < 10^-20^ for all comparisons) as well as an enrichment of pathogenic mutations on residues with significantly high G scores (Supplementary Fig. 1e-f). As the pathogenicity of mutations in ClinVar is all supported by clinical or experimental evidence^3^, these results provide virtual clinical and experimental validation of the efficacy of using known disease mutation-derived G scores as a predictive feature.

**Figure 2.**
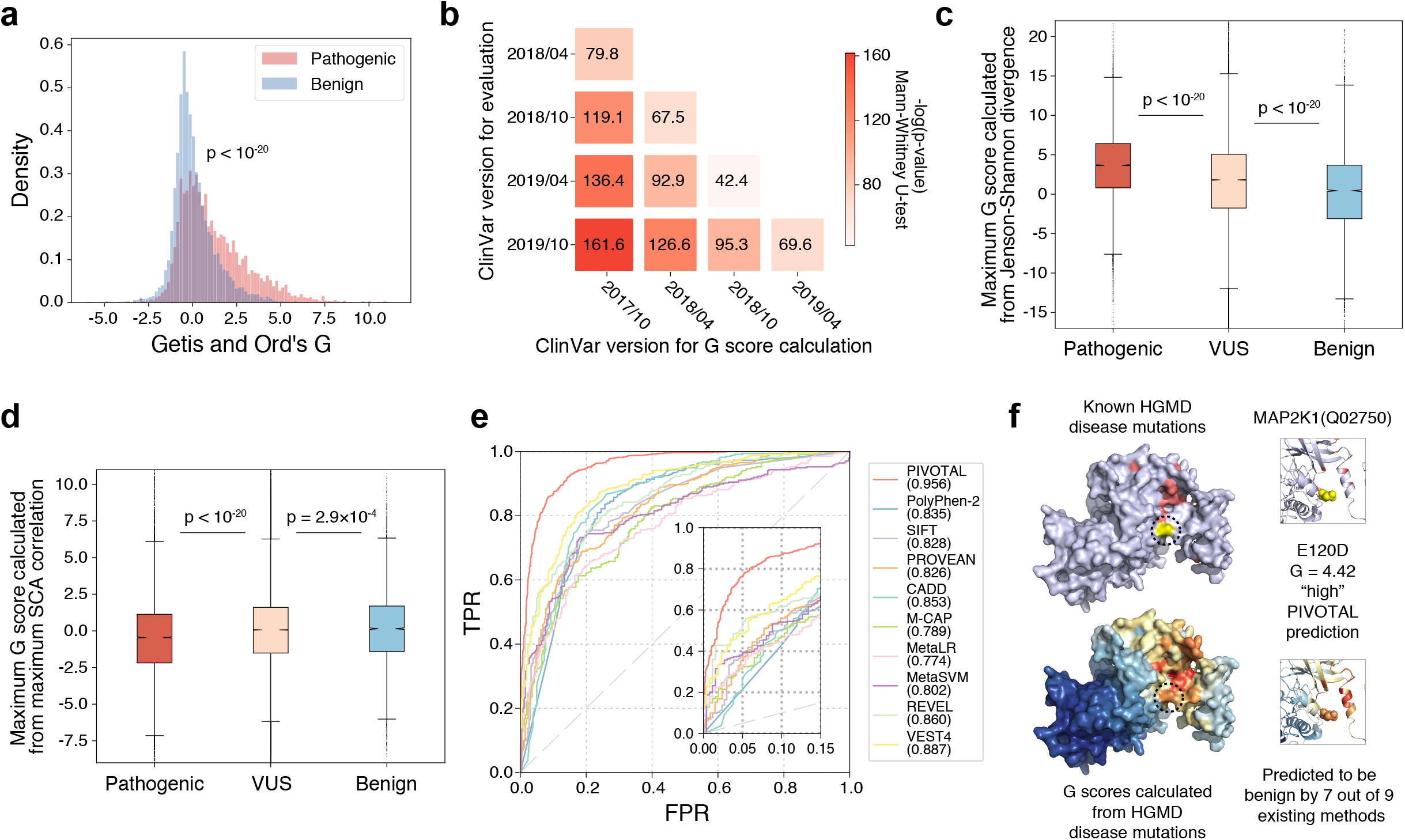
The general G scores and the combined PITOVAL score for VUS prioritization. (a) ClinVar pathogenic mutations have significantly higher G scores than benign mutations (P < 10^-20^, Mann-Whitney U test) when the absence or presence of known disease mutations in HGMD is used for G score calculation. (b) ClinVar pathogenic mutations have significantly higher G scores than benign mutations when the absence or presence of known pathogenic mutations in a previous version of ClinVar is used for G score calculation, irrespective of the versions used for G score calculation and evaluation. (c) G scores calculated from Jenson-Shannon divergence are significantly higher for pathogenic mutations than for VUS mutations (P < 10^-20^, Mann-Whitney U test), which in turn have significantly higher scores than benign mutations (P < 10^-20^, Mann-Whitney U test). (d) G scores calculated from maximum intra-protein SCA correlation serve to significantly separate pathogenic, VUS and benign mutations (P < 10^-20^ for both comparisons, Mann-Whitney U test), with pathogenic mutations having lower scores than benign mutations. (e) The combined PITOVAL score outperforms existing pathogenicity predictors on the left-out test set in classifying ClinVar pathogenic and benign mutations, and this advantage is especially significant when the critical region where the false positive rate is low. (f) An example of a known pathogenic mutation, E120D, in the *MAP2K1* gene, predicted to be pathogenic with “high” confidence respectively by PIVOTAL, whilst 7 out of 9 existing pathogenicity predictors predicted it to be benign. Its close spatial proximity to a number of known HGMD disease mutations indicated by its high disease-specific G scores is one of the main reasons for its successful prediction by PIVOTAL.

Evolutionary conservation has been an important feature for existing pathogenicity predictors^11–14,27^. Indeed, we show that ClinVar pathogenic mutations are significantly more conserved than VUS (P < 10^-20^, Mann-Whitney U test), which in turn have higher conservation scores than benign mutations (Supplementary Fig. 2a, P < 10^-20^, Mann-Whitney U test). However, a survey of current computational tools has discovered that most of them falsely predict a high proportion of benign variants at highly conserved positions as pathogenic and often fail to predict truly pathogenic variants at less conserved positions^27^. The G statistic addresses this weakness by considering the conservation of the overall structural neighborhood of a residue. Indeed, G scores calculated from conservation exhibit significant separation between pathogenic, VUS and benign mutations (Fig. 2c, P < 10^-20^ for both comparisons, Mann-Whitney U test), and that a high G score does not require high conservation at the residue itself (Supplementary Fig. 2b). In addition to conservation, sequence co-variation has also been shown to yield substantial predictive power^28,29^. By examining the maximum intra-protein correlation for each residue with statistical coupling analysis (SCA), we find that pathogenic mutations tend to occur at amino acid residues that co-evolve less with other residues (Supplementary Fig. 2c, P < 10^-20^, Mann-Whitney U test), and that this can only be partially attributed to the fact that these residues are highly conserved, as shown by the slight negative correlation between JS divergence and maximum SCA correlation (Supplementary Fig. 2d). Under a similar rationale with evolutionary conservation, G statistics calculated from coevolution significantly differ between mutations of different clinical significance (Fig. 2d, P < 10^-20^ for both comparisons, Mann-Whitney U test) while providing complementary information to SCA correlation itself (Supplementary Fig. 2e). Notably, these G scores, conservation and co-evolution features do not exhibit high correlation with predictions from existing variant prioritization tools on either the dataset comprised of known pathogenic and benign mutations (Supplementary Fig. 2f) or the VUS dataset (Supplementary Fig. 2g), whereas prediction scores from existing methods show much closer correlation. This further demonstrates that our G score features, conservation, and co-evolution provide orthogonal information to existing variant pathogenicity prediction methods.

To test whether these features confer additional power upon existing variant prioritization tools, we combined G scores calculated from known disease mutations, conservation and co-evolution, as well as raw conservation and maximum co-evolution scores, with existing pathogenicity predictors (PolyPhen-2, SIFT, PROVEAN, CADD, M-CAP, MetaLR, MetaSVM, REVEL and VEST4). We discovered that a combined XGBoost model, trained by incorporating the three types of G scores mentioned above and raw conservation and co-evolution scores, rendered significantly higher performance than the existing predictor alone, regardless of the predictor of choice (Supplementary Fig. 3a-i). These results suggest that integrating these novel metrics, derived from spatial distribution patterns of known disease mutations and genomic properties, provide complementary predictive power that allows us to better prioritize VUS. The 9 existing pathogenicity predictors of our choice represent the state-of-the-art and most commonly used ones, and their source of predictive power covers all important features used for pathogenicity prediction, namely, features derived from protein structures, protein sequence conservation, DNA-level genomic conservation as well as genomic and epigenomic annotations.

Next, we constructed an XGBoost classifier with all three types of G scores, JS divergence, maximum SCA correlation, as well as all 9 existing pathogenicity predictors mentioned above. We notice that G scores calculated from known disease mutations and conservation could serve as pathogenicity classifiers alone with fair performance, while G scores calculated from co-evolution alone is a weak classifier (Supplementary Fig. 4a). The final PIVOTAL classifier reaches an area under the receiver operating characteristic curve (AUROC) of 0.956, substantially outperforming any single existing predictor, especially in the critical region with a small false positive rate (Fig. 2e). More interestingly, the final classifier exhibits substantial improvement over a classifier trained using only the 9 existing pathogenicity predictors as features (Supplementary Fig. 4b-c), further manifesting the efficacy of incorporating additional information including 3D spatial neighborhood (G scores), conservation and coevolution. As an example for demonstrating the improvement of PIVOTAL over existing tools, a pathogenic mutation in the test set, E120D in MAP2K1^30^, are predicted by PIVOTAL as having “high” potential (see Methods) to be pathogenic, yet 7 out of 9 existing tools predicted it to be a benign mutation. A homology model of MAP2K1 shows that both mutations are in close proximity to a number of known disease mutations, as characterized by their high G statistics calculated from known disease mutations (Fig. 2f). Notably, this mutation also has a significantly high G score for JS divergence (Supplementary Fig. 4d) but not a significantly low G score for co-evolution (Supplementary Fig. 4e), indicating the significant contribution of the clustering of known disease mutations as well as conserved residues in its structural neighborhood to our final prediction. Using this final PIVOTAL classifier, we calculated prediction scores for all 143,293 VUS mutations in the latest ClinVar release (Supplementary Table 2).

Variants in the same gene can be associated with multiple clinically distinct disorders^31^. However, for VUS in these pleiotropic genes, there is currently no tool to predict the specific disease with which they are potentially associated, despite efforts to predict their molecular mechanisms of pathogenicity^32^. To this end, we developed a disease-specific G score by tailoring the disease mutation-based G score such that only known pathogenic mutations associated with a specific disease are used for calculation. Since disease terms are inter-connected and the same disease phenotype can be described at different levels of the ontology of disease terms^33^ (Fig. 3a), we constructed a directed acyclic graph (DAG) for disease terms annotated for pathogenic variants (see Methods), and calculated disease-specific G scores for all terms across different levels of the ontology. As anticipated, G scores for the annotated disease terms of pathogenic variants are significantly more elevated than other disease terms on the corresponding 3D protein structures (Supplementary Fig. 5a). As disease terms at higher levels may be annotated for more mutations, thereby rendering more information-rich G statistics, for each pathogenic variant in the latest ClinVar release, we separated all disease terms into two categories: ancestors (including itself), which are more general descriptions of the annotated disease term for that variant, and other disease terms, which the variant is not associated with (Supplementary Fig. 5b). We found that G scores for ancestors of the annotated disease terms of pathogenic variants are, as expected, significantly higher than those for other disease terms (Fig. 3b). Moreover, residues harboring pathogenic mutations are much more likely to exhibit a significantly high G score for ancestors of the annotated disease terms than for other disease terms across all FDR cutoffs (Fig. 3c). These results indicate that the relative ranking of disease-specific G scores for different terms is suggestive of the actual associated disease. On the other hand, for VUS and benign mutations annotated with a disease term, disease-specific G scores for the term itself as well as its ancestor terms are significantly higher for VUS mutations than benign mutations (Fig. 3d), further confirming the validity of this disease-specific G score as an indicator of pathogenic potential regarding a specific disease phenotype.

**Figure 3.**
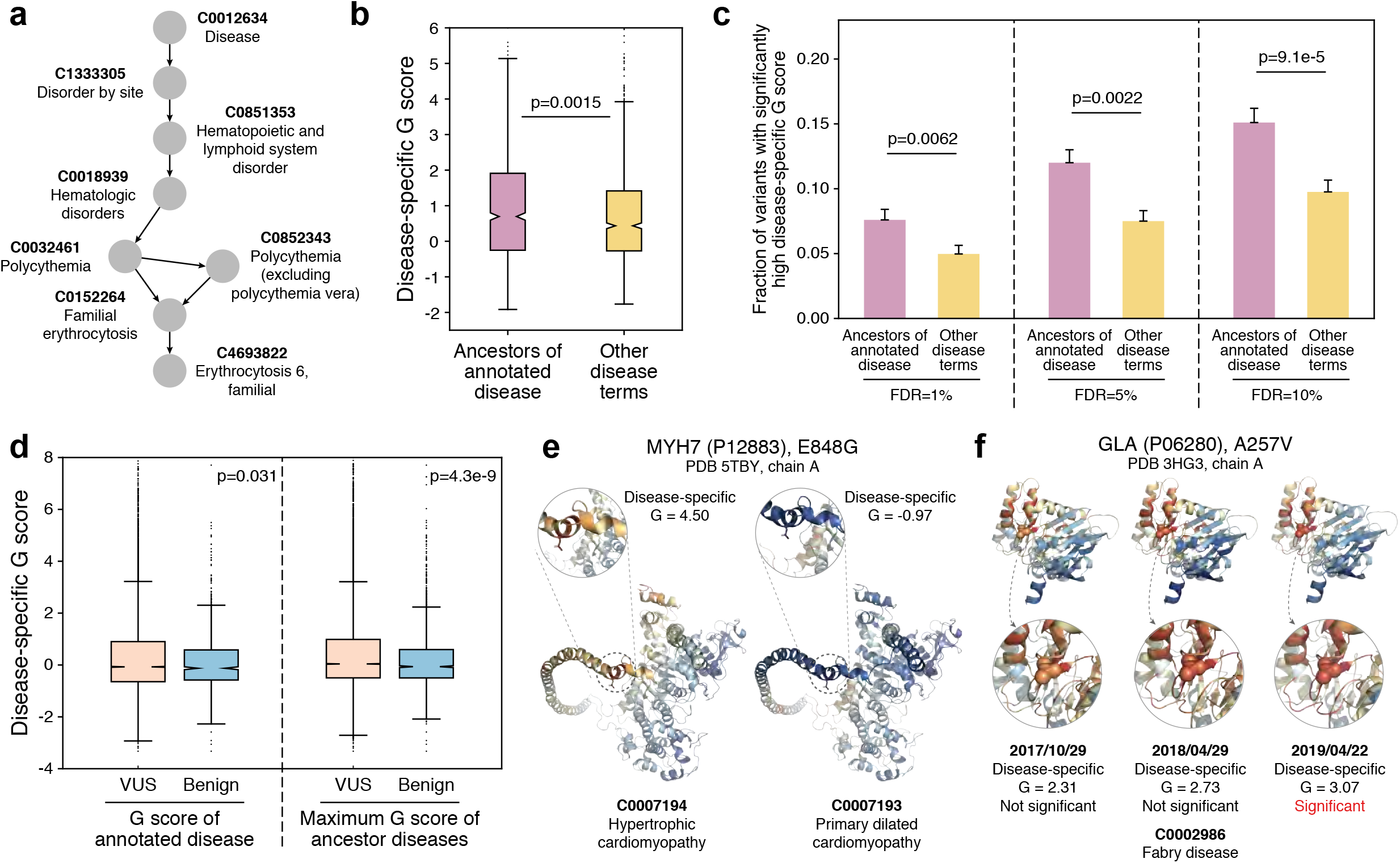
The disease-specific G score for disease association inference. (a) An example of the disease term hierarchy. (b) Disease-specific G scores for ancestors of the annotated disease term of pathogenic variants are significantly higher than those for other disease terms (P = 0.0015, Mann-Whitney U test). (c) Residues harboring pathogenic mutations have a significantly higher chance of having a significantly high G score for ancestors of the annotated disease term than for other disease terms across all FDR cutoffs (P-values calculated using the Mann-Whitney U test). (d) For VUS and benign mutations annotated with a phenotype term, disease-specific G scores for the term itself (P = 0.031, Mann-Whitney U test) as well as its ancestor terms (P = 4.3×10^-9^, Mann-Whitney U test) are significantly higher for VUS mutations than benign mutations. (e) An example of a known disease mutation in the *MYH7* gene, E484G, causing hypertrophic cardiomyopathy and has no known association with dilated cardiomyopathy. Disease-specific G score of this mutation for hypertrophic cardiomyopathy is significantly elevated whereas that for dilated cardiomyopathy is not. (f) An example of a mutation on the *GLA* gene, A257V, known to cause Fabry disease. The disease-specific G score of this mutation for Fabry disease increased from 2.31 in 2017, which was not significantly high, to 3.07 in early 2019, which was significantly elevated, demonstrating the steady increase in power of PIVOTAL as more information becomes available.

As an example to demonstrate the efficacy of this disease-specific G score in pinpointing the potential disease association of variants, *MYH7* is a pleiotropic gene known to be associated with two clinically distinct cardiomyopathies: hypertrophic cardiomyopathy (HCM) and dilated cardiomyopathy (DCM)^34^. A missense mutation, E848G, has a high G score of 4.50 for HCM yet a low G score of −0.97 for DCM (Fig. 3e). This pathogenic mutation has been detected in HCM patients from multiple sources^35,36^ and has been reported to disrupt myofibril contraction^37^ possibly by disrupting the proteinprotein interaction of MYH7 with cardiac myosin binding protein C (cMyBP-C)^36^, yet it is not known to be associated with DCM, which usually involves reduced functions of the myosin motor domain^36,38^. Although previous studies have implicated the differential distributions of HCM and DCM mutations on the protein structure of MYH7^38^, PIVOTAL goes one step further and provides a means of quantifying such distributions to give clinically meaningful evaluations about the potential disease association of variants.

It is worth noticing that the power of PIVOTAL increases through time as the clinical significance of more variants become available. To illustrate this, a missense mutation, A257V, on the *GLA* gene which encodes alpha-galactosidase A, has been reported to reduce galactosidase activity by 46%^39^ and is classified as a likely pathogenic variant for Fabry disease in ClinVar. Using only pathogenic mutations causing Fabry disease reported in a late 2017 version of ClinVar, the disease-specific G score of this mutation for Fabry disease is 2.31, which fails to achieve statistical significance at 5% FDR after multiple testing corrections. However, as more disease annotations became available for GLA variants, the disease-specific G score of this mutation increased to 2.73 in early 2018, and further reached 3.07 with an early 2019 version of ClinVar (Fig. 3f), where residue 257 was identified as having a significantly high G score for Fabry disease. As more disease associations of variants become available in the future, our PIVOTAL framework will have even more capability to not only prioritize VUS but also highlight specific diseases that they are potentially associated with.

To facilitate the exploration of the spatial distribution of human missense variation and genomic properties in the context of the structural proteome, we have built the PIVOTAL web server (Fig. 4) where users can visualize clinically detected variants, evolutionary conservation and co-evolution for all 17,605 human proteins with at least one structural model, as well as obtain general G statistics calculated from all three types of information, disease-specific G scores, in addition to output from the final classifier which allows users to prioritize variants of their interest. Users can also search by disease terms to find all proteins harboring mutations causing a specific disease or calculate G statistics for custom user-provided genome properties with its “G score calculator” functionality. With future increases to the scale of clinical variation databases and the structural coverage of the human proteome, we expect PIVOTAL to provide even more insightful information for both clinicians attempting to evaluate the pathogenic potential and possible disease associations of VUS and scientists performing functional research on missense variation.

**Figure 4.**
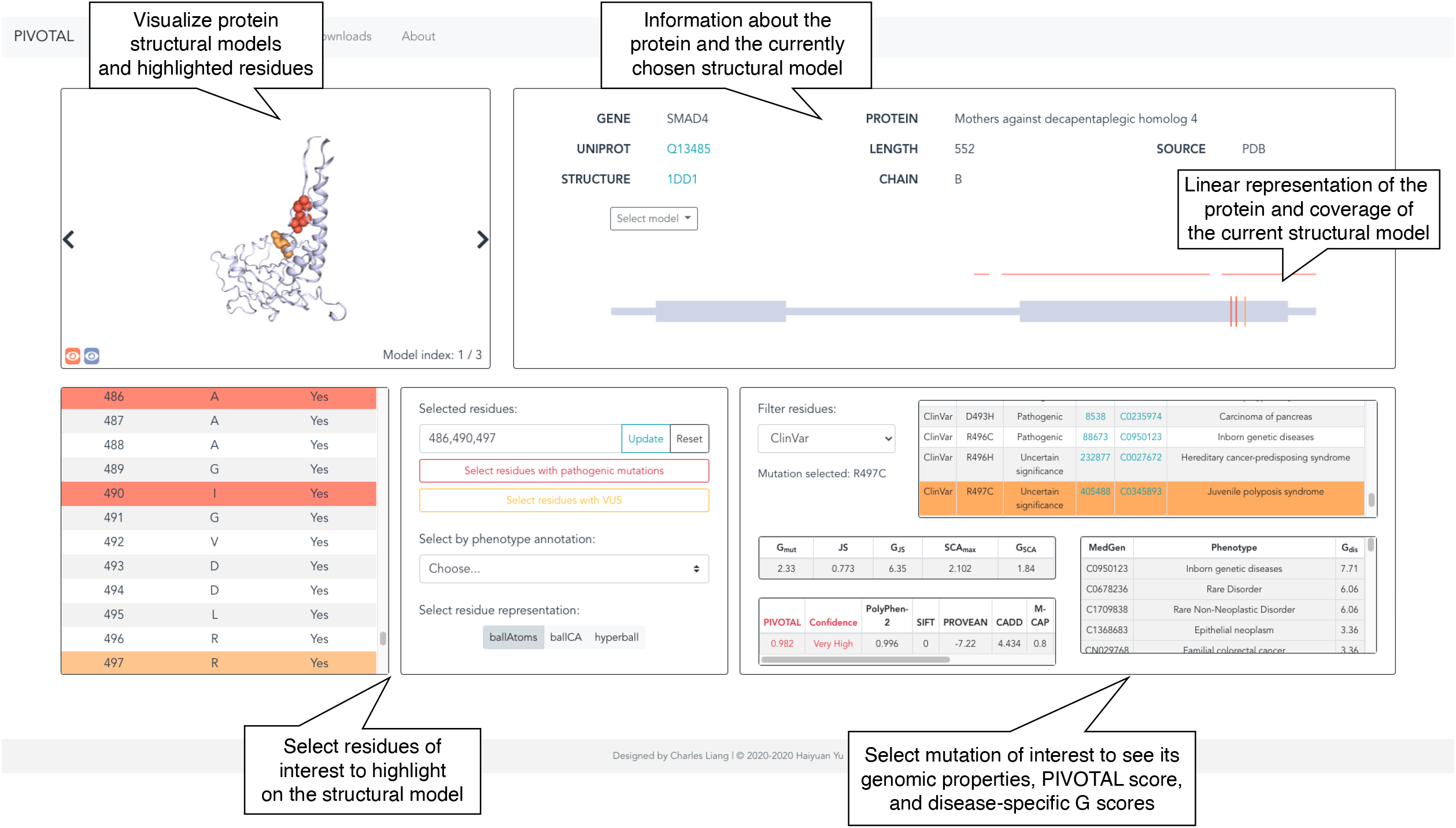
The PIVOTAL web tool. This figure depicts a result page after a user has queried a protein or gene of interest where they can visualize protein structural models with selected residues highlighted, and obtain information about the protein, the structural models, as well as variants in the gene along with PIVOTAL predictions and disease-specific G scores for VUS.

## METHODS

### Construction of the human 3D structural proteome

To build a comprehensive repository of structural models for all human proteins, we collected experimentally determined structures from the PDB as well as homology models from ModBase^40^ for all canonical isoforms of human Swiss-Prot entries obtained from UniProt. For PDB chains, we collected residue mappings between PDB and UniProt entries from SIFTS^41^, and for homology models we only retained those having an MPQS score of at least 0.5. We filtered out structural models that covered less than 10% of the full length of the protein, and for each protein we sorted all available models first by source (PDB preferred over ModBase) and then by coverage (high coverage preferred). To remove redundancy, for each protein we started from the model with the highest priority and included an additional model only if it covered 5 additional amino acid residues unseen by models that were already included. Finally, our human 3D structural proteome comprised 7,185 PDB chains and 23,532 ModBase homology models for a total of 17,605 proteins (Supplementary Table 3).

### Calculation of Getis and Ord’s local G statistic

For each structural model, Getis and Ord’s local G statistics^24,25^ were calculated from a residue-level raw score (e.g. presence or absence of disease mutations, evolutionary conservation, or co-evolution) ***x***, and a spatial weight matrix ***W*** that defines the spatial relationship between each pair of residues. For disease mutation-based G scores, residues harboring known disease mutations have a raw score of 1, otherwise the raw score is zero. Here we used an inverse distance weight method where the weight between two two different residues *i* and *j* is inversely proportional to their distance (in Å): 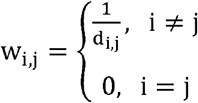, where *d_i,j_* is the distance between the Cα atoms of residues *i* and *j*. The G score for each residue *i* is then calculated as:

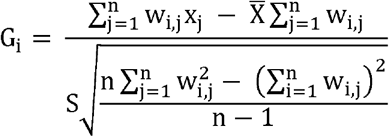

where *X_j_* is the raw score for residue *j, n* is the number of residues in the structural model, and

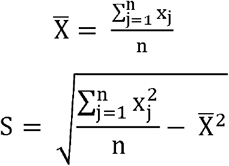

The G statistic is a z-score (standard score) from which a p-value can be calculated. Residues with significantly high or low raw scores were identified by comparing the adjusted p values to the significance level after Benjamini-Hochberg correction.

The G statistic was calculated on structural models having at least 30 amino acid residues. In order to obtain a single G score for each residue of a protein, we aggregated G scores calculated from different structural models of the same protein by taking the maximum. For the identification of residues with significantly high G scores on a protein, we took the union of residues identified from different structural models.

### Compilation of pathogenic, benign and VUS mutations

We compiled known pathogenic missense mutations from two databases: the Human Gene Mutation Database (HGMD, version 2019/02) and ClinVar. We collected all disease-causing missense mutations (DM) from HGMD and 82,920 of them were mapped to UniProt identifiers and residues through the Ensembl Variant Effect Predictor (VEP)^42^. Five ClinVar versions were collected with a timestamp interval of about half a year, the latest being 2019/10/21. To obtain sets of missense variants that could be confidently identified as pathogenic, benign or VUS at each timestamp, we obtained submission records at the corresponding timestamp. Our pathogenic variants comprised variants whose overall clinical significance is “pathogenic” or “likely pathogenic”, have at least one submission classifying them as “pathogenic” or “likely pathogenic”, and have no submission classifying them into other categories. Similarly, our benign variants consisted of variants whose overall clinical significance is “benign” or “likely benign”, have at least one submission classifying them as “benign” or “likely benign”, and have no submission classifying them into another category. On the other hand, our VUS variants comprised those that are assigned an overall clinical significance of “uncertain significance”, have at least one submission classifying them as “uncertain significance”, and have no submission classifying them as “pathogenic” or “benign”. The latest ClinVar version we collected had 30,494 pathogenic variants, 17,818 benign variants and 143,293 VUS. Statistics for other timestamps can be found in Supplementary Table 1.

### Calculation of evolutionary conservation and intra-protein co-evolution

To generate multiple sequence alignments (MSAs) for all human proteins, we ran PSI-BLAST^43^ for all human Swiss-Prot protein sequences against all protein sequences in UniProt downloaded in June 2019 to search for homologous sequences. Only eukaryote protein sequences with an PSI-BLAST e-value below 0.05 covering at least 50% but not identical to the query protein were retained, and we imposed a limit of one best hit per species. MSAs were subsequently generated with Clustal Omega^44^ with default parameters, and the output was trimmed to remove any position in the MSA where there was a gap in the query protein. From these MSAs, Jenson-Shannon divergence^45^ was calculated with a custom Python script as a measure of evolutionary conservation. Intra-protein co-evolution was calculated by statistical coupling analysis (SCA)^46^, which outputs a correlation matrix where the rows and columns each correspond to all residues. To generate a residue-level raw score for G score calculation, we took the maximum SCA correlation for each residue as it provides us with information on the maximum strength of co-evolution with another residue.

### Construction of a classifier for VUS prioritization

To construct a classifier for prioritizing VUS by incorporating G scores with existing pathogenicity prediction tools, we collected 10,035 ClinVar pathogenic or benign missense variants (as filtered by our criteria stated above) on 3,357 proteins in the 2019/10/21 version on residues that do not harbor HGMD (version 2019/02) missense DM mutations. All variants were split into a training set and a test set by splitting all proteins such that these two sets do not have variants on the same protein. The resulting training set had 7,528 variants, whereas the validation and test set had 2,507 variants, respectively. For each variant, G scores calculated from disease mutation, JS divergence and maximum SCA correlation, as well as raw JS divergence and maximum SCA correlation scores, were used as features, in addition to 9 existing variant effect predictors: PolyPhen-2, SIFT, PROVEAN, CADD, M-CAP, MetaLR, MetaSVM, REVEL and VEST4. We fitted an XGBoost classifier with training data and tuned its hyperparameters with 5-fold cross-validation by maximizing the area under the receiver operating characteristic curve (AUROC) of the validation set using random search. Note that when splitting the training data into different folds, we made certain that no two folds have variants on the same protein. After hyperparameter optimization, the classifier was re-fitted with the entire training set and evaluated on the leave-out test set, which was unseen during classifier fitting and tuning. The final classifier was re-fitted on the entire training and test data and used for making predictions. Predictions were provided as both the raw prediction score from the classifier as well as a tiered prediction confidence constituting *Very High*, *High*, *Medium, Low* and *Very Low,* whose cutoff were determined by quintiles of all raw prediction scores.

### Curation of disease terms

Both ClinVar and HGMD provide annotations of variant phenotypes in MedGen Concept IDs. However, since the same disease can be annotated by multiple disease terms in different levels of generality (for example, see Fig. 3a), variants causing the same disease can have different disease term annotations. To address this problem, we obtained relationships between MedGen^47^ concepts from MGREL.RRF available from the MedGen FTP site and extracted all “CHD” (child) and “PAR” (parent) relations between two different terms. In cases where two terms are both a parent/child of the other, the two terms were merged. A directed acyclic graph (DAG) was then constructed from all terms with parent/child relationships and the largest connected component was retained. To further simplify the graph, we started from a “Disease” root term (C0012634) and included new terms (nodes) and relationships (edges) in a layer-wise fashion. Starting from the root term as the first layer, nodes in each new layer must be previously undiscovered and connected to a node in the previous layer. A set of disease terms annotated for over 20 pathogenic mutations yet not included in this simplified DAG were manually curated and added to the DAG. This resulting simplified DAG was used for all analyses of the disease-specific G score.

### PIVOTAL web interface

To build the PIVOTAL web interface, we leveraged the handiness and flexibility of the Django web framework as well as a number of popular JavaScript packages, including JQuery, Bootstrap, in addition to D3, which is used for drawing two-dimensional geometric figures. The interactive display of 3D protein structures is empowered by the NGL^48^ and MichelaNGLo^49^ JavaScript packages. Our modifications to these packages enable users to select a specific structural model for a protein, highlight specific residues on both the 2D representation and the 3D structure of the protein simultaneously, and switch between different visual representations.

---

**Supplementary Figure 1.** Summary statistics and enrichment of pathogenic mutations on residues with significantly high HGMD disease mutation-derived G scores. (a) Type distribution of ClinVar coding variants. (b) Fraction of pleiotropic and non-pleiotropic genes for genes harboring at least one HGMD disease (DM) mutation. (c) Fraction of pleiotropic and non-pleiotropic genes for genes harboring at least one ClinVar pathogenic or likely pathogenic mutation. (d) ClinVar pathogenic mutations are enriched in residues with significantly elevated G scores calculated from HGMD disease mutations at all FDR cutoffs (*: P < 0.001, P values calculated from two-sided Z-tests). (e) ClinVar pathogenic mutations from a later version are enriched in residues with significantly elevated G scores calculated from ClinVar pathogenic mutations from an earlier version, irrespective of the versions of choice. Odd ratios are displayed. (f) Negative log P values of the log odds ratios in (e), indicating significant enrichment for all version combinations (P-values calculated from two-sided Z tests).

**Supplementary Figure 2.** Raw evolutionary conservation and co-evolution scores provide additional power for distinguishing pathogenic and benign variants. (a) Pathogenic mutations are significantly more likely to occur on evolutionarily conserved residues than VUS (P < 10^-20^, Mann-Whitney U test), which occur on significantly more conserved residues than benign mutations (P < 10^-20^, Mann-Whitney U test). (b) An example of a known pathogenic mutation, R113H in the *EIF2B5* gene, that does not occur in an evolutionarily conserved residue with a low JS divergence but nevertheless has a significantly high G score calculated from JS divergence. (c) Pathogenic mutations are significantly more likely to occur on residues that co-evolve less with other residues than VUS (P < 10^-20^, Mann-Whitney U test), which occur on residues co-evolving significantly less with other residues than benign mutations (P = 9.5×10^-6^, Mann-Whitney U test). (d) JS divergence and maximum SCA correlation are only slightly negatively correlated, indicating their distinct contribution to the performance of the PIVOTAL classifier. (e) An example of a known pathogenic mutation, M331R in the *STAT3* gene, that has a high maximum SCA correlation score yet has a significantly low G score calculated from SCA correlation, indicating the different information conferred by the raw coevolution score and the G score. (f) Pairwise Pearson correlation between features on the training dataset. (g) Pairwise Pearson correlation between features on the VUS dataset.

**Supplementary Figure 3.** Performance comparison between an XGBoost classifier constructed from new features (three types of G scores, JS divergence and SCA correlation) along with an existing pathogenicity classifier and the existing classifier alone, the existing classifier being (a) PolyPhen-2, (b) SIFT, (c) PROVEAN, (d) CADD, (e) M-CAP, (f) MetaLR, (g) MetaSVM, (h) REVEL, (i) VEST-4. The combined classifier outperforms the existing classifier regardless of the tool of choice.

**Supplementary Figure 4.** Performance of the PIVOTAL classifier. (a) Receiver operating characteristic (ROC) curves using each G score feature alone to predict pathogenicity. (b) Receiver operating characteristic (ROC) curves comparing PIVOTAL with an XGBoost meta-classifier combining all 9 existing pathogenicity predictors. (c) Precision-recall curves comparing PIVOTAL with an XGBoost meta-classifier combining all 9 existing pathogenicity predictors. (d) The same mutation as shown in Fig. 2f, E120D in *MAP2K1,* with its JS divergence and G score calculated from JS divergence (structure colored by G scores calculated from JS divergence, with blue indicating low values and red indicating high values). (e) The same mutation as shown in Fig. 2f, E120D in *MAP2K1,* with its maximum SCA correlation and G score calculated from SCA correlation (structure colored by G scores calculated from SCA correlation, with blue indicating low values and red indicating high values).

**Supplementary Figure 5.** The disease-specific G score. (a) Disease-specific G scores for the annotated disease term of pathogenic variants are distributed significantly more highly than those of other disease terms (P = 0.023, Mann-Whitney U test). (b) For every variant annotated with a disease term, all disease terms in the constructed directed acyclic graph can be divided into two categories: ancestors of the annotated disease term (labeled in purple), which all describe the annotated disease in different levels of generality, and other terms (labeled in yellow), which do not describe the annotated disease term. This division forms the basis of comparison in Fig. 3b and Fig. 3c.

**Supplementary Table 1.** Summary statistics of the five ClinVar versions collected.

**Supplementary Table 2.** PIVOTAL predictions for all 143,293 ClinVar VUS mutations.

**Supplementary Table 3.** Information of all 30,717 structural models used in PIVOTAL.

