## Supplementary figures and images for "PIVOTAL: Prioritizing variants of uncertain significance with spatial genomic patterns in the 3D proteome"

### Supplementary_Figure_1.pdf

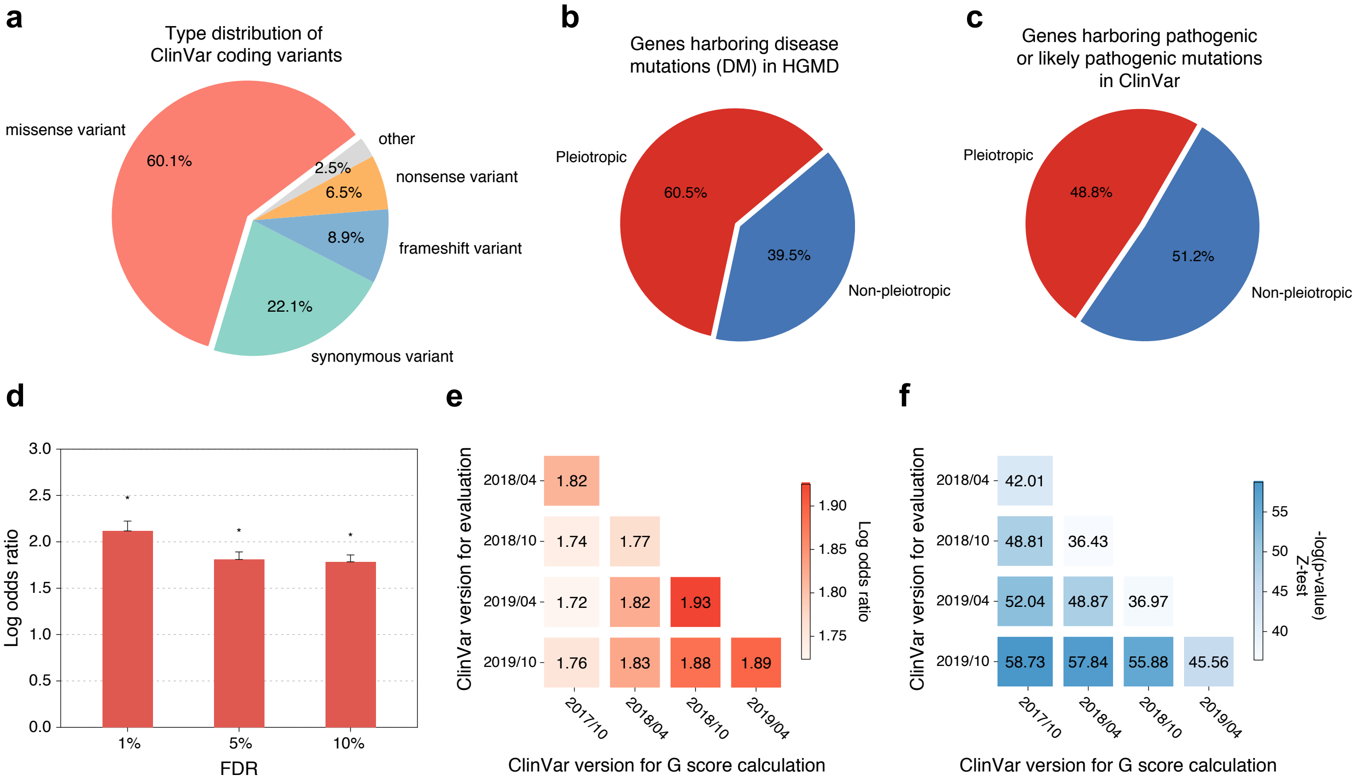

### Supplementary_Figure_2.pdf

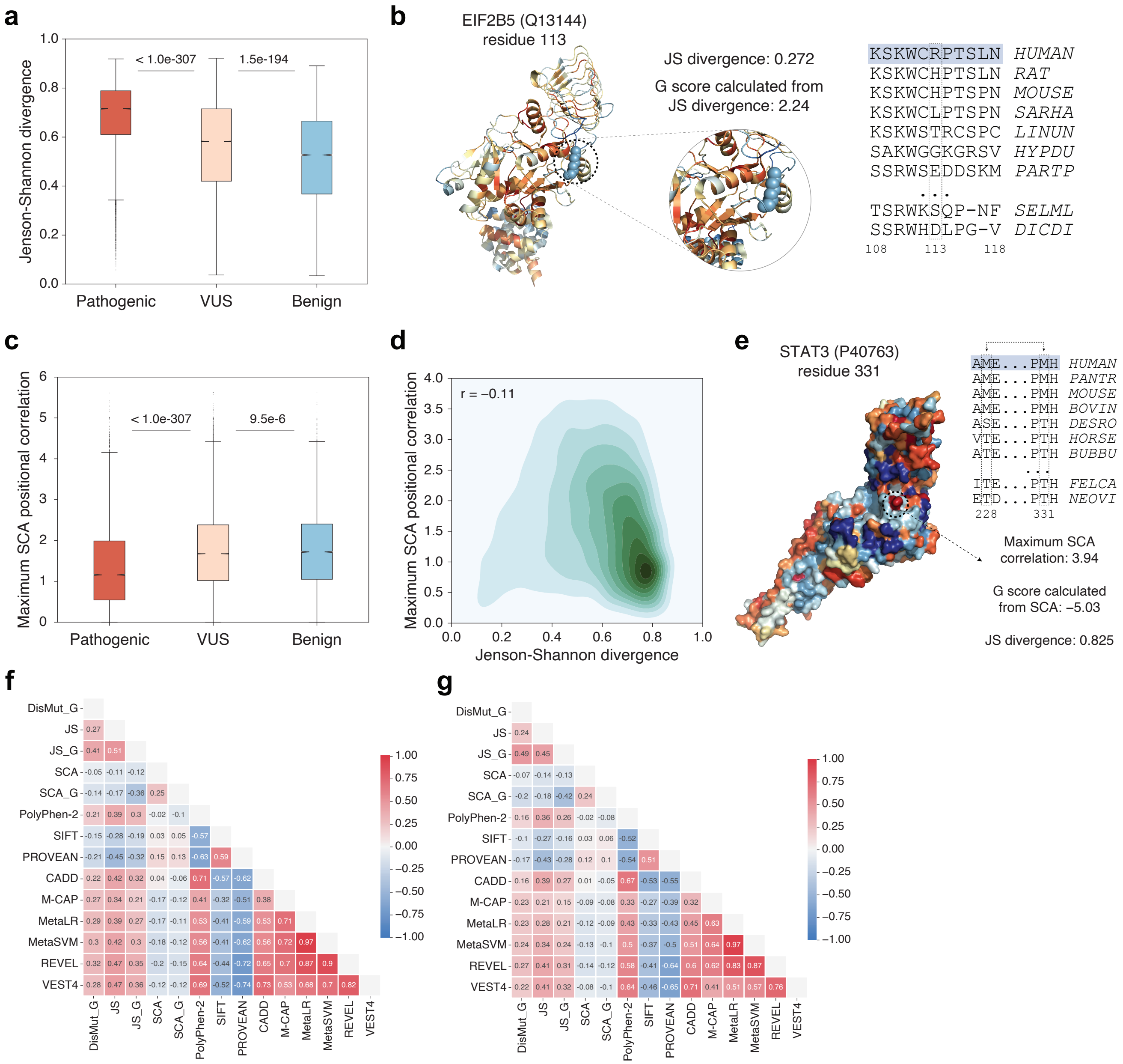

### Supplementary_Figure_3.pdf

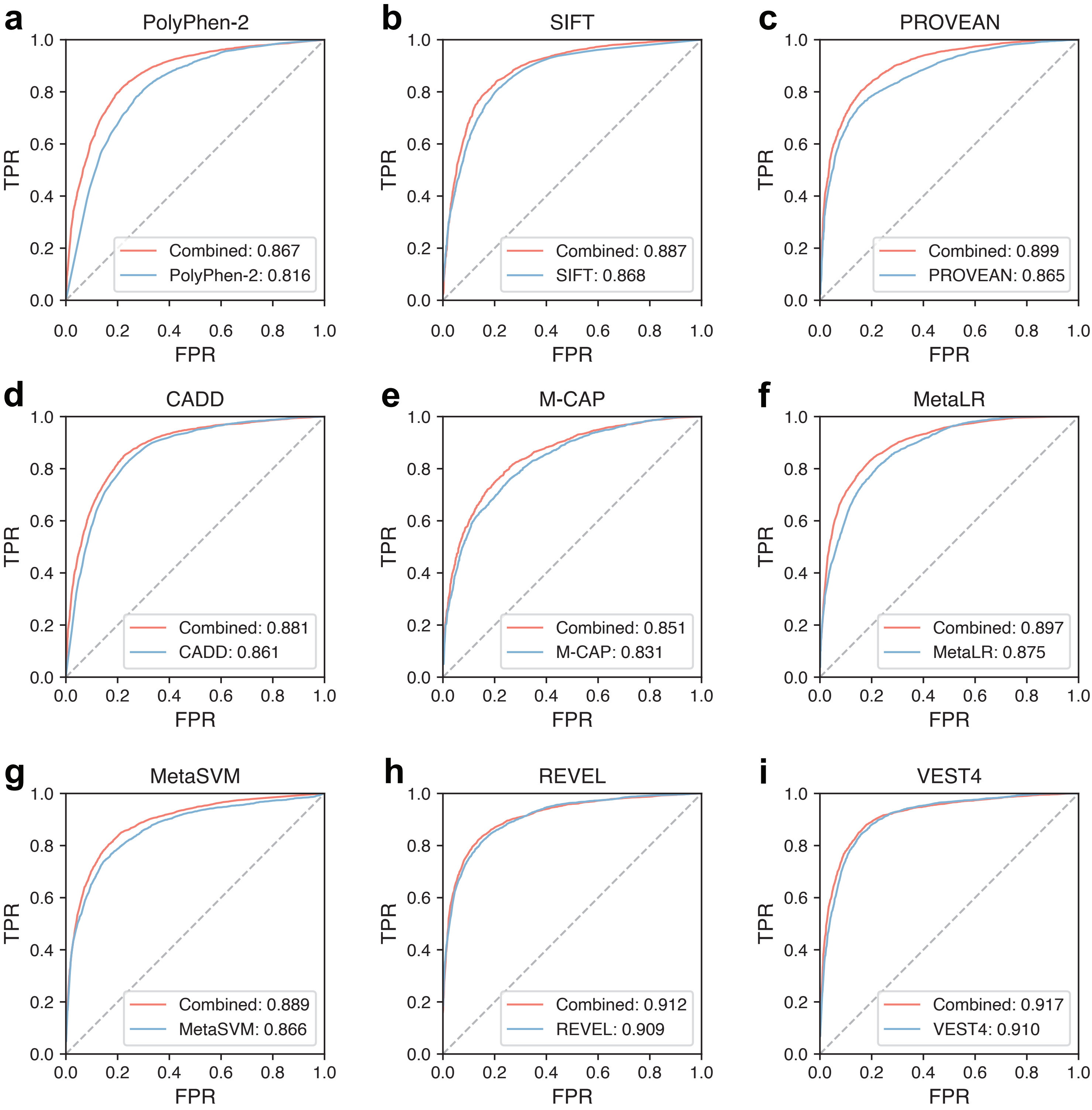

### Supplementary_Figure_4.pdf

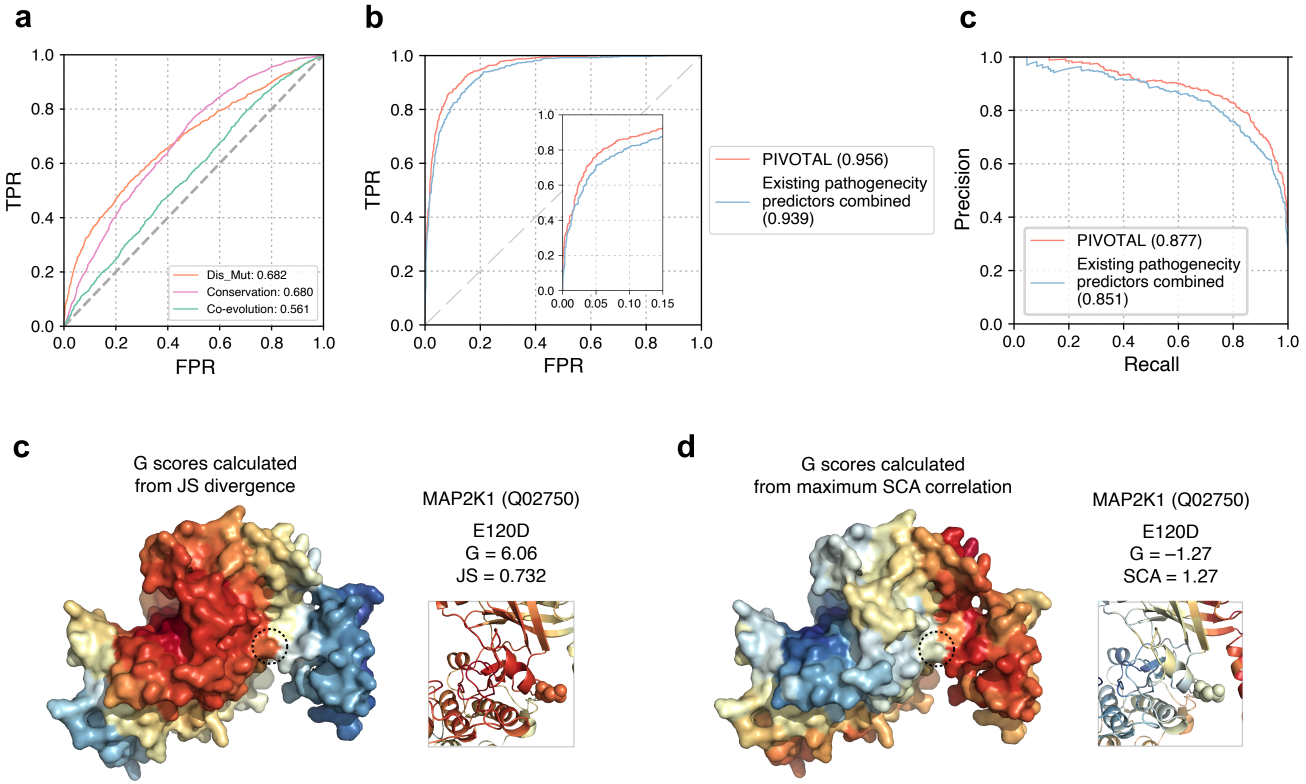

### Supplementary_Figure_5.pdf

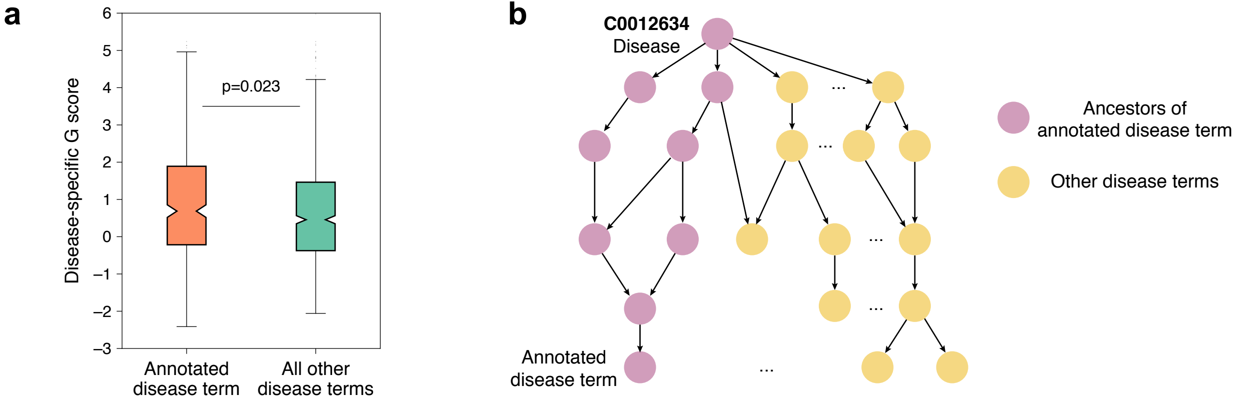
